# Inhibition of advanced glycation end product formation and serum protein infiltration in bioprosthetic heart valve leaflets: Investigations of anti-glycation agents

**DOI:** 10.1101/2022.02.28.482351

**Authors:** Andrey Zakharchenko, Christopher A. Rock, Tina Thomas, Samuel Keeney, Emily Hall, Hajime Takano, Abba M. Krieger, Giovanni Ferrari, Robert J. Levy

## Abstract

Bioprosthetic heart valves (BHV) fabricated from glutaraldehyde pretreated heterograft tissue, such as bovine pericardium (BP) are the most commonly used heart valve replacements. However, BHV durability is limited by structural valve degeneration (SVD) resulting from both calcification and advanced glycation end product (AGE) deposition together with serum protein infiltration. In the present study we investigated the hypothesis that anti-AGE agents, Aminoguanidine (AG), Pyridoxamine (PYR), and N-Acetylcysteine (NAC) could mitigate AGE-serum protein mechanisms in model studies, both in vitro and in vivo, using rat subdermal implants of BP. In vitro studies demonstrated that each of these agents could significantly inhibit AGE formation in BP. However, in rat 28 days BP subdermal implants, only PYR demonstrated both significant inhibition of AGE and serum albumin accumulation per immunostaining. BHV calcification was not mitigated by PYR. It is concluded that AGE-serum protein pathophysiology contributing to SVD can be ameliorated by PYR.

## 1. Introduction

Heart valve disease (HVD) affects millions worldwide and yet there is no medical therapy for it. Ultimately HVD is approached by surgical valve replacement. Valve prostheses, such as mechanical or bioprosthetic heart valves (BHV), are commonly used to treat patients with severe symptomatic HVD. While mechanical valves offer extended durability compared to BHV, their use is associated with increased thrombosis risk, requiring lifelong anticoagulation therapy, with associated risks of hemorrhage. On the other hand, BHV offer a better hemodynamic profile and in most cases do not require lifelong anticoagulation. Furthermore, recent advances in BHV technology enable a minimally invasive transcatheter delivery approach with substantial reduction in mortality rate and improved quality of life[1, 2].

Despite these benefits, BHV are subject to calcification and structural valve degeneration (SVD), ultimately resulting in valve failure. While calcification was thought to be a major culprit for SVD, recent studies suggest the important role of the inflammatory processes within BHV. Importantly, it was documented that patients with diabetes would have a significantly reduced BHV lifetime due to advanced glycation end product (AGE) formation within leaflets. Protein glycation and AGE formation are the hallmarks of diabetic pathophysiology leading to collagen crosslinking, tissue stiffening, and formation of pro-inflammatory epitopes. Recent studies by our group have shown formation of AGE including both AGE crosslinks and pro-inflammatory AGE epitopes in clinical BHV explants[3]. Furthermore, BHV incubations in vitro in accelerated AGE conditions results in disruption of mechanical and hydrodynamic properties. Collectively these data provide evidence that AGE accumulation in BHV is a major risk factor for SVD that needs to be addressed.

To address the severe effects of AGE in diabetes, experimental drug candidates were explored to mitigate AGE formation. Among the most studied candidates were aminoguanidine (AG), Pyridoxamine (PYR), and N-acetyl cysteine (NAC), which were shown to have significant antiglycation effects both in vivo and in vitro and were involved in multiple clinical trials. Studies have shown that AG, PYR, and NAC act as sacrificial scavengers, reacting with AGE intermediates and reducing AGE formation [4, 5]. Additionally, NAC interacts with the inflammatory receptor for AGE (RAGE) resulting in its downregulation[6]. Systematic administration of AG in diabetic rats resulted in mitigation of AGE-related diabetic complications, such as cataract formation, collagen tissue glycation, and oxidative species formation. Administration of PYR resulted in reduction of levels of circulating AGE intermediates, such as glyoxal (GLX), and reactive oxygen species in diabetic rats. It was shown that during an in vitro incubation of human bone tissue under hyperglycemic conditions, AG acted as a scavenging agent, reacting with AGE intermediates. This resulted in significant reduction of glycation in comparison to the control[7]. To explore diabetic vascular complications in vivo, a diabetic rat model was used where AG was shown to have an effect on glycation and reactive oxygen species production. Collectively, these data provide the rationale for investigating anti-glycation effects of AG, PYR, and NAC towards mitigation of AGE in BHV.

The current study tested the hypothesis that AGE inhibitors (AG, PYR, NAC) would significantly reduce BP AGE formation in established in vitro and in vivo glycation models. Glutaraldehyde-fixed bovine pericardium (BP), the most commonly utilized heterograft material to fabricate BHV leaflets, was used to model pathophysiological BHV processes of AGE formation and calcification. Incubation of BP with AGE precursors, with or without AGE inhibitors, was monitored for 28 days. Collagen structure, radiolabeled glucose uptake, and collagen shrink temperature were monitored to assess the degree of structural degeneration. In vivo studies involved a well-established juvenile rat subdermal model to study glycation and calcification. Following BP implantation, AGE inhibitors were administered for 7 or 28 days. Following sacrifice, BP explants were analyzed for mechanistic and structural changes including serum albumin uptake, AGE formation, and collagen structure change. Blood levels of AGE metabolites and biomarkers were monitored and compared with the levels in control unoperated animals of the same age. Thus, this work provides novel insights into the potential use of AGE inhibitors for patients with HVD.

## 2. Materials and Methods

### 2.1 Materials

#### 2.1.2 Glutaraldehyde Fixation of Bovine Pericardium

Fresh BP purchased from Animal Technologies (Tyler, TX) was rinsed with saline (0.9% NaCl) and fixed in a 0.6% glutaraldehyde solution in 50 mM HEPES pH 7.4 buffer (Sigma 51558) for 7 days at room temperature. After 7 days, the BP was transferred into a 0.2% glutaraldehyde storage solution at 4°C. Prior to each experiment, BP was rinsed in phosphate-buffered saline (PBS) buffer (Sigma 806552) to remove unbound glutaraldehyde.

#### 2.1.3 Sample Preparation for In Vitro Studies

Glutaraldehyde-fixed BP samples (n=5) were punched using 8 mm biopsy punches (Medline, Northfield, IL). Samples were rinsed with PBS and then incubated for 28 days in PBS with the following conditions either without inhibitors (Control) or with inhibitors (n-acetylcysteine (NAC, 5mM), pyridoxamine (PYR, 50mM), or aminoguanidine (AG, 10mM): PBS only, glucose (GLU, 100 mM) only, glucose (100mM) + Bovine serum albumin (BSA) (5%) (GLU-BSA), or BSA (5%) only. Incubation solutions contained sodium azide (0.1%) to maintain sterility. After 28 days, samples were exhaustively rinsed with PBS prior to further analysis.

#### 2.1.4 Rat Subdermal Implants

Male juvenile (3 weeks old, average weight 60 grams) Sprague Dawley rats (Charles-River Laboratories, Wilmington, MA) received subdermal BP implants under general anesthesia, as previously described. All protocols used were approved by the Institutional Animal Care and Use Committee at The Children’s Hospital of Philadelphia. Each animal was implanted with two 1 cm^2^ glutaraldehyde-fixed BP patches, with a total of 10 rats per time point and 20 rats per treatment group. For the treatment groups, the animals received either NAC (500 mg/mL), AG (500 mg/mL), or PYR (200 mg/mL) via drinking water. For the control groups, the animals received subdermal implants but had no drugs in the drinking water. The animals were sacrificed 7 days or 28 days post-implantation. The BP explants were rinsed with sterile saline. Each explant was cut into four pieces, with two pieces stored in 10% neutral buffered formalin for histological processing and two pieces frozen for calcium content analysis.

#### 2.1.5 Radiolabeled Glycation Assays

To quantify the glycation kinetics of the model substrates, samples (n=5 per group) were punched from BP using an 8 mm biopsy punch. These samples were incubated for 1-28 days at 37°C shaking at 110 RPM in the 100 mM solution of glucose supplemented with 0.544μCi/ml ^14^C-labeled glucose with or without inhibitors (NAC, PYR, or AG). All solutions contained sodium azide (0.1%) to maintain sterility. The radioactivity of each medium was measured at the start of the incubation to calibrate the radioactive signal and at the end of the incubation to confirm sufficient excess of radioactive reagents (<10% drop in radioactive signal). This was done by taking a 500μl aliquot, adding 5 mL of Bioscint, shaking the solution till it turned clear, and then performing scintillation counting. Scintillation counting was performed on a Beckman LS 6000 (Beckman Coulter; Brea, CA).

Following incubations, the samples were extensively rinsed with deionized water and lyophilized for at least 48 hours. The dried samples were weighed and then dissolved using 500 μl Biosol per sample and incubating at 50°C until fully dissolved. To measure the radioactivity of each sample, 5 mL of Bioscint was added and the solution was shaken till turning clear before performing scintillation counting.

The scintillation count was used to calculate glucose incorporation into the substrate normalized by dry weight. To study the effects of inhibitors on the BSA glycation, 5% BSA was added to the 100 mM glucose solution supplemented with 0.544μCi/ml ^14^C-labeled glucose with or without inhibitors (NAC, PYR, or AG) for 1, 3, 7, 14, or 28 days at 37°C shaking at 110 RPM and 1 mL aliquots were taken at each timepoint. Glycated BSA was separated from unreacted glucose using gravity-flow chromatography columns (Bio-Rad 7322010) and protein concentration was measured with BCA assay (Sigma 71285-M). Then, 500 μl aliquots were taken and 5 ml of Bioscint was added and the solution was shaken until it turned clear before performing scintillation counting. The scintillation count was used to calculate glucose binding BSA molecules with or without AGE inhibitors.

#### 2.1.6 Differential Scanning Calorimetry Analysis

To determine the effect of glycation conditions with AGE-inhibitors on collagen crosslinking in vitro and in vivo, differential scanning calorimetry (DSC) was used. A DSC7 (PerkinElmer) instrument was used to determine collagen denaturation temperature, as previously described. In brief, both in vitro BP samples (n=5) and in vivo BP explants (n=10) were punched into circles using a 4mm biopsy punch. Samples were then blotted on the paper to remove excess liquid, placed in an aluminum pan and sealed. Sealed samples were placed into a calorimeter with temperature ramping from 25°C to 100°C.

### 2.2 Morphological Studies

#### 2.2.1 Two-Photon Microscopy

To visualize collagen microarchitecture, two-photon microscopy was used. For in vitro and in vivo samples, either BP punches or BP explants were mounted onto a silicon chamber (Electron Microscopy Sciences, Hatfield, PA) and covered with PBS in between two glass coverslips. Second harmonic generation (SHG) imaging was performed using a custom Prairie Technologies Ultima multiphoton system (Middleton, WI) attached to an Olympus BX-61 upright microscope (Tokyo, Japan). The system contains GaAsP photomultiplier tubes and a diode-pumped broadband mode-locked titanium: sapphire femtosecond MaiTai HP laser (Spectra Physics, Irvine, California). Imaging was done by focusing the laser on the sample using a 40X water-immersion objective (Olympus xlumpfln, Tokyo, Japan). Scans were performed using an excitation wavelength of 920 nm with a 505 nm short pass filter. Images were analyzed to measure the collagen crimp period using custom MATLAB (Mathworks; Natick, MA) scripts, as previously described [8].

#### 2.2.2 Immunohistochemical Staining

Tissue designated for IHC was fixed in 10% neutral buffered formalin at 4°C for 48 hours. The tissue was then gradually dehydrated and embedded in paraffin. The paraffin blocks were sectioned at 6 μm and mounted on Histobond (VWR; Radnor, PA) slides. The slides were then heated in an oven and re-hydrated in successive xylene to ethanol baths. The slides were then incubated overnight in 60°C citrate buffer (Thermo Fisher; Waltham, MA) for antigen retrieval. Following antigen retrieval, slides were rinsed then incubated with primary antibody. At the following day slides were blocked using a goat serum (Abcam ab7481) to minimize non-specific binding. Next, slides were incubated with primary antibodies (αAGE, 400 ng/ml| ab23722; αBSA, 38 ng/ml|ab192603) overnight at 4°C. After incubation with primary antibodies, slides were washed and incubated with 3% hydrogen peroxide for 10 minutes. Slides were then rinsed and incubated for 1 hour at room temperature with the appropriate horseradish peroxidase polymer-conjugated secondary antibody (Abcam; Cambridge, United Kingdom). Slides were then rinsed and incubated for 8 minutes at room temperature with 3,3’Diaminobenzidine substrate (Abcam; Cambridge, United Kingdom). Slides were then counter-stained using regressive hematoxylin staining, dehydrated, and cover-slipped.

#### 2.2.3 Von Kossa Staining

For visualization of calcium deposits on rat explants, Von Kossa Staining was applied for both 7-day and 28-day rat explants according to manufacturer’s protocol (Abcam ab150687).

### 2.3 Quantitative Assays

#### 2.3.1 Calcium Analysis

For calcium quantification, 7-day and 28-day rat explants were lyophilized, weighed, and hydrolyzed with 6M hydrochloric acid. Hydrolyzed samples were then analyzed using a colorimetric assay according to manufacturer’s protocol (Sigma MAK022).

#### 2.3.2 Enzyme Immunosorbent Assays

Quantification of levels of AGE, methylglyoxal, and methylglyoxal protein adducts were done using ELISA in accordance with manufacturer’s instructions. During the in vivo studies, serum samples were collected from each rat and were analyzed according to the following kits: methylglyoxal (Abcam ab238543), RAGE (Abcam ab202409), Methylglyoxal assay kit (Abcam ab241006).

### 2.4 Statistical Methods

For crimp period and shrink temperature analyses to determine the significance of the changes for each incubation compared to the PBS control Dunnett’s test was used to control for multiplicity. Statistical analyses of IHC images’ intensity were performed using 1-way ANOVA and treatments were compared to the control. For all statistics tests, p<0.05 was considered significant. In cases where there might be concern that the data are skewed and hence P-values using conventional normal theory might be suspect, we also performed nonparametric tests: where the two-sample t-tests was performed a Wilcoxon rank sum test was run; where a one-way analysis of variance was performed a Kruskal-Wallis test was run. The resulting p-values for the nonparametric tests were substantively the same.

## 3. Results

### 3.1. Effect of glycation and AGE inhibitors on the BP collagen structure

For second-harmonic generation microscopy (SHG) studies (Figure 1), the integrity of BP collagen structure was shown to be disrupted by glycation and serum protein infiltration. Accumulation of AGE within the BHV results in collagen crosslinking, formation of pro-inflammatory epitopes, and the alteration of mechanical properties [8]. To study the effects of glycation and AGE inhibitors, BP samples were exposed to AGE precursors (BSA, GLU, and GLU-BSA) with or without AGE inhibitors (AG, NAC, PYR). The changes in collagen structure were monitored using SHG, a non-invasive optical approach that allows for imaging inner collagen structures. Unmodified BP had a well-organized collagen structure with distinct crimp bands (Fig. 1A). Co-incubation of BP with GLU-BSA for 28 days resulted in collagen disorganization (Fig. 1B) and crimp period increase (Fig. 1D), while co-incubation in the presence of AGE inhibitors preserved collagen structure (Fig. 1C). The effect on collagen morphology disruption was less pronounced after a separate incubation of BP with either GLU or BSA (Supplementary Fig. 1) suggesting the essential role of glycated albumin in SVD. Quantitative analysis of crimp period showed no significant changes in crimp distance after 28 days of incubation in PBS (Fig. 1D) or any of the PBS-inhibitor co-incubations, suggesting minimal interactions between AGE inhibitors and BP collagen (Fig. 1D, PBS). BP incubation with BSA for 28 days resulted in no significant changes in average crimp value (Fig. 1D, BSA). Co-incubations with AG or PYR did not result in significant changes in crimp period, compared to the 28-day PBS control, yielding crimp periods of 14.92±0.1 μm and 13.96 ±1.18 μm, respectively; however, incubation with NAC resulted in a significant increase in crimp period (16.86±1.2 μm) (Fig. 1D, BSA).

**Figure 1.**
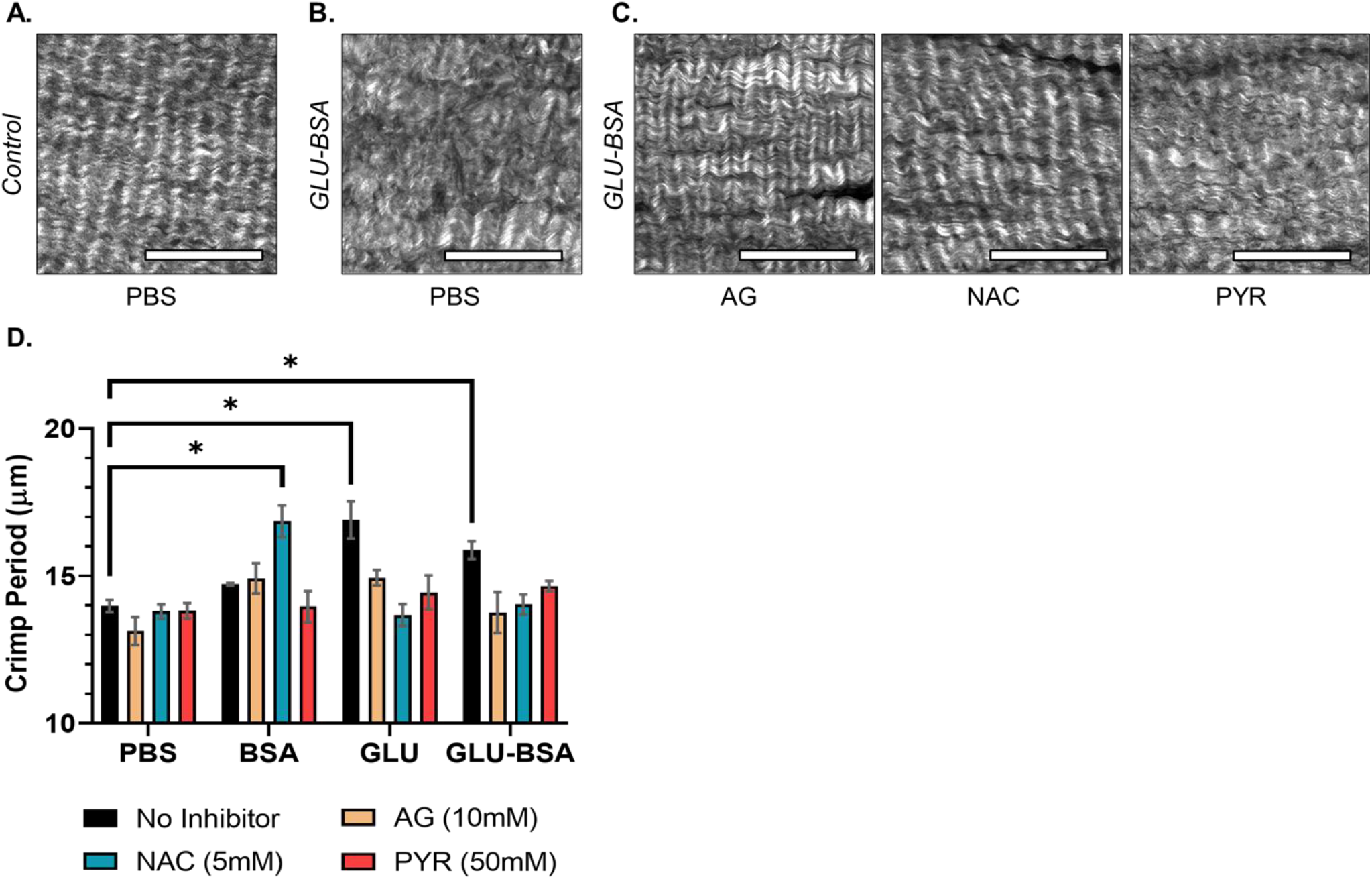
Two-photon microscopy images and crimp period image analysis of glutaraldehyde-fixed bovine pericardium (BP) before and after 28-day exposure to glycation inhibitors: none (PBS), 10mM aminoguanidine (AG), 5mM n-acetylcysteine (NAC), and 50mM pyridoxamine (PYR). **A:** Representative image of BP in PBS prior to incubation. **B:** BP after co-incubation with 100 mM glucose and 5% BSA (GLU-BSA) for 28 days without inhibitors (PBS). **C:** Representative images of BP co-incubated with GLU-BSA and inhibitors. Scale bars are 100 μm. **D:** Crimp period analyses of BP incubated with inhibitors and 1) PBS, 2) PBS with 5% bovine serum albumin (BSA), 3) PBS with 100 mM glucose (GLU), 4) PBS with 100 mM glucose and 5% BSA (GLU-BSA). Error bars indicate standard error. Significance was determined Dunnett’s Method, using 28-day PBS as a control (*p < 0.05).

Incubation with 100 mM glucose solution for 28 days resulted in a significant increase in collagen crimp to 16.9±1.41 μm, which was reduced by all three of the inhibitors (14.94±0.58 μm for AG, 13.68±0.81 μm for NAC, and 14.44±1.29 μm for PYR). Co-incubation of GLU-BSA with BP also resulted in a significant increase in crimp period compared to control (15.88±0.67 μm), which was mitigated by all of the inhibitors: AG (13.76±1.55 μm), NAC (14.04±0.76 μm), and PYR (14.66 ±0.39 μm) (Fig. 1D, GLU-BSA).

Collagen denaturation temperature, or shrink temperature (ST), as an index of crosslinking, was measured after incubation with AGE inhibitors following SHG imaging to assess the effects of inhibitors and glycation on degree of collagen crosslinking. There was no significant change in ST after 28-day incubation in PBS (88±0.4°C) or NAC (87.4±0.5°C), compared to the 0-day control (87.8±0.4°C) (Fig. 2A). Co-incubation with PYR (86.5±0.2°C) or AG (85.7±0.3°C) resulted in a significant drop in ST (Fig. 2A). Co-incubation with BSA did not result in a change of ST in control (88.2±0.4°C) or PYR (87.3±0.4°C), while both NAC and AG resulted in a significant drop with 86.4±0.4°C and 85.9±0.3°C, respectively (Fig. 2B). Glucose co-incubation resulted in a significant drop in ST in all of the inhibitor groups: 85.6±0.38°C in AG, 85.8±0.3°C in NAC, and 85.4±0.4°C in PYR (Fig. 2C). This data was consistent with the crimp period mitigation effect observed in SHG (Fig. 1D, GLU).

**Figure 2.**
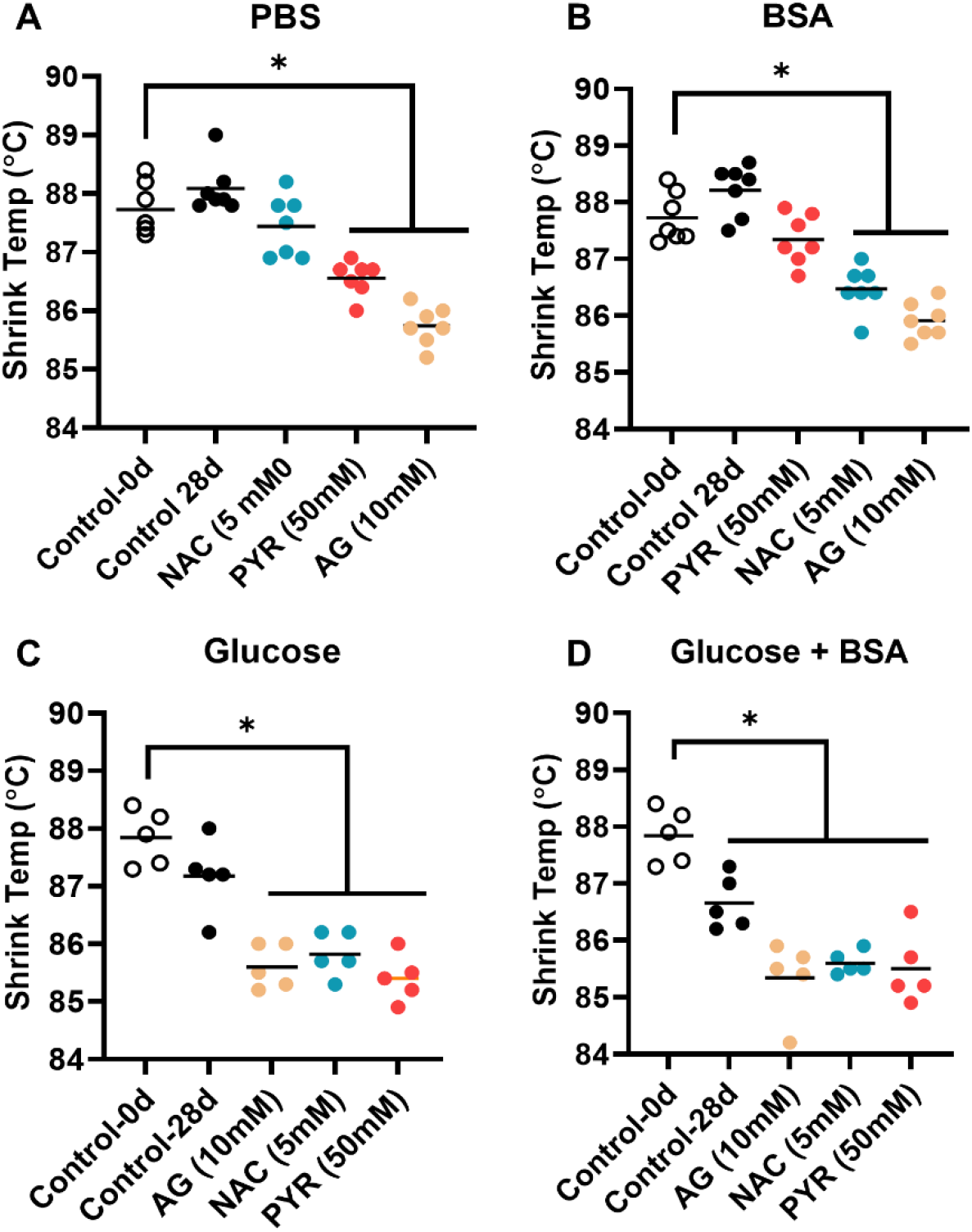
Shrink temperature of glutaraldehyde-fixed BP as determined by differential scanning calorimetry before and after 28 days of exposure to the following glycation inhibitors: prior to incubation (Control-0d), none (Control-28d), aminoguanidine (AG 10mM), n-acetylcysteine (NAC, 5mM), and pyridoxamine (PYR, 50mM). **A.** Inhibitors in PBS. Pyridoxamine and aminoguanidine showed significant (*p < 0.05) decrease in shrink temperature. **B.** Inhibitors in PBS with 5% bovine serum albumin (BSA). NAC and AG showed significant (*p < 0.05) decrease in shrink temperature. **C.** Inhibitors in PBS with glucose (100mM). All three inhibitors showed significant (*p < 0.05) decrease in shrink temperature. **D.** Inhibitors in PBS with glucose (100mM) plus BSA (5%). All three inhibitors showed significant decrease in shrink temperature. Significance was determined by Dunnett’s method comparing to no inhibitor. (*p < 0.05).

No change was detected in the glucose-incubated control (87.1±0.6°C) (Fig. 2C). Combining BSA and GLU resulted in a significant reduction in all groups: 86.6±0.4°C in control, 85.3±0.6°C in AG, 85.6±0.2°C in NAC, and 85.5±0.6°C in PYR (Fig. 2D). These results suggest that interaction between AGE inhibitors and BP tissue does not change macroscopic collagen structure.

### 3.2 Effect of AGE inhibitors on glucose incorporation

In order to understand the nature of molecular interactions between BP and AGE inhibitors under glycation, ^14^C-labeled glucose radioactive studies were done. These experiments sought to study the effects of AGE inhibitors on kinetics and extent of glucose incorporation into BP or BSA, respectively. The rationale of these studies was based on the established BP and BSA glycation models[8]. BSA glycation was studied as BSA is the most abundant serum protein found in blood and BHV explants and it is highly susceptible to glycation[9]. BP was incubated with ^14^C radiolabeled glucose for 1-28 days with or without inhibitors and the amount of permanently bound glucose was measured.

The glucose uptake by control BP demonstrated exponential growth during the first 14 days that plateaued at 28 days with 10.7±1.1 nmol/mg, similar to previously published data (Fig. 3A) [8]. All three inhibitors had a significant mitigating effect on the glucose uptake after 28 days with roughly 50% inhibition by AG and PYR (4.9±0.5 nmol/mg and 5.0 ±1.2 nmol/mg respectively), whereas NAC demonstrated only 38% inhibition with 6.7±0.9 nmol/mg (Fig. 3A).

**Figure 3.**
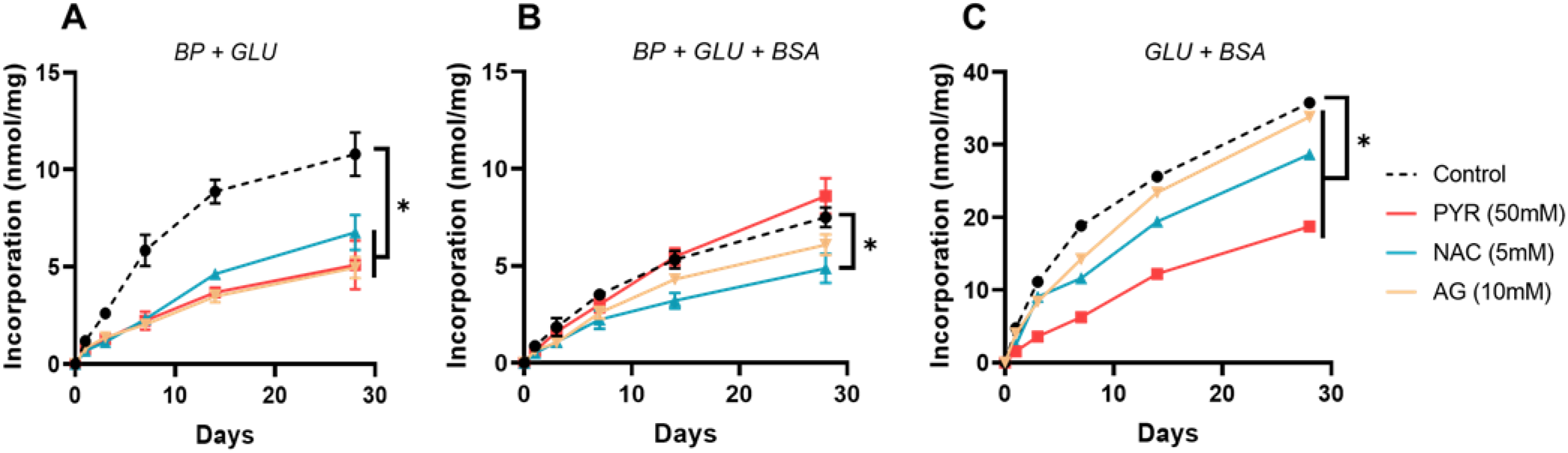
Glycation kinetics of glutaraldehyde-fixed bovine pericardium (A, B) or bovine serum albumin (BSA, 5%) over 28 days when exposed to ^14^C-labeled glucose along with the following glycation inhibitors: none (Control), pyridoxamine (PYR, 50mM), aminoguanidine (AG, 10mM), and n-acetylcysteine (NAC, 5mM). **A.** Incorporation when exposed to glucose (100mM). All three inhibitors showed significant decreases (*p < 0.05) in glucose incorporation from 24 hours onward. **B.** Incorporation when exposed to glucose (100mM) plus bovine serum albumin (BSA, 5%). NAC showed significant decrease (*p < 0.05) in glucose incorporation from 24 hours onward. **C.** All three inhibitors showed significant decreases (*p < 0.05) in glucose incorporation from 24 hours onward. Data shown are means of 5 replicates. Error bars indicate standard deviation. Significance was determined by Dunnett’s method comparing the control to no inhibitor at each time point.

Co-incubation of BP with BSA-GLU resulted in a nearly 50% decrease of glucose uptake by the tissue with 5.69±1.06 nmol/mg (Fig. 3B). There was a significant mitigation effect from NAC (4.86±0.75 nmol/mg) and no effect from AG or PYR, 6.0±0.5 nmol/mg and 8.59±0.9 nmol/mg, respectively (Fig. 3B). This could be due to a competitive reaction between BP collagen and BSA with glucose in solution. While there are a large number of accessible amino groups in BSA, the majority of BP collagen amino groups are functionalized with glutaraldehyde and therefore do not participate in reaction. At the same time, BP is highly susceptible to BSA infiltration, as was shown earlier, therefore two processes are taking place: BSA uptake by BP and BSA glycation by glucose.

To understand the effects of AGE inhibitors on BSA glycation, radiolabeled glucose was incubated with 5% BSA with or without inhibitors (Fig. 3C). 28-day glucose uptake by BSA was 330% higher than by BP alone (Fig. 3B control vs. 3A control) and there was a significant inhibition by NAC and PYR, with 28.67±0.4 nmol/mg and 18.76±0.2 nmol/mg, respectively (Fig. 3C). By contrast, AG demonstrated minimal inhibition after 28 days with 28.67±0.4 nmol/mg glucose uptake (Fig. 3C). Thus, BSA was much more susceptible to glycation in solution, which could be mitigated by AG and PYR.

### 3.3 Rat subdermal implant studies: comparisons between AGE inhibitors in vivo

The rat subdermal implantation model using juvenile rats is a well-established approach used by many groups to study BP treatments on calcification [10]. Furthermore, clinical comparisons demonstrated that subdermal implantation adequately models albumin uptake and glycation of BP tissue, similar to what is seen in BHV clinical explants [3, 11]. BP samples were implanted in juvenile rats and AGE inhibitors were administered. Samples were explanted at 7 or 28 days and characterized in terms of collagen structure, ST, albumin infiltration, AGE formation, and calcification (Fig. 4). BP examination by SHG revealed noticeable collagen crimp disorganization in 7 and 28 days compared to the unimplanted control (Fig. 4A). There was a noticeable improvement by all of the inhibitors in all 7 and 28 days, however the changes were not significant after quantitative analysis (Fig. 4B, 4C).

**Figure 4.**
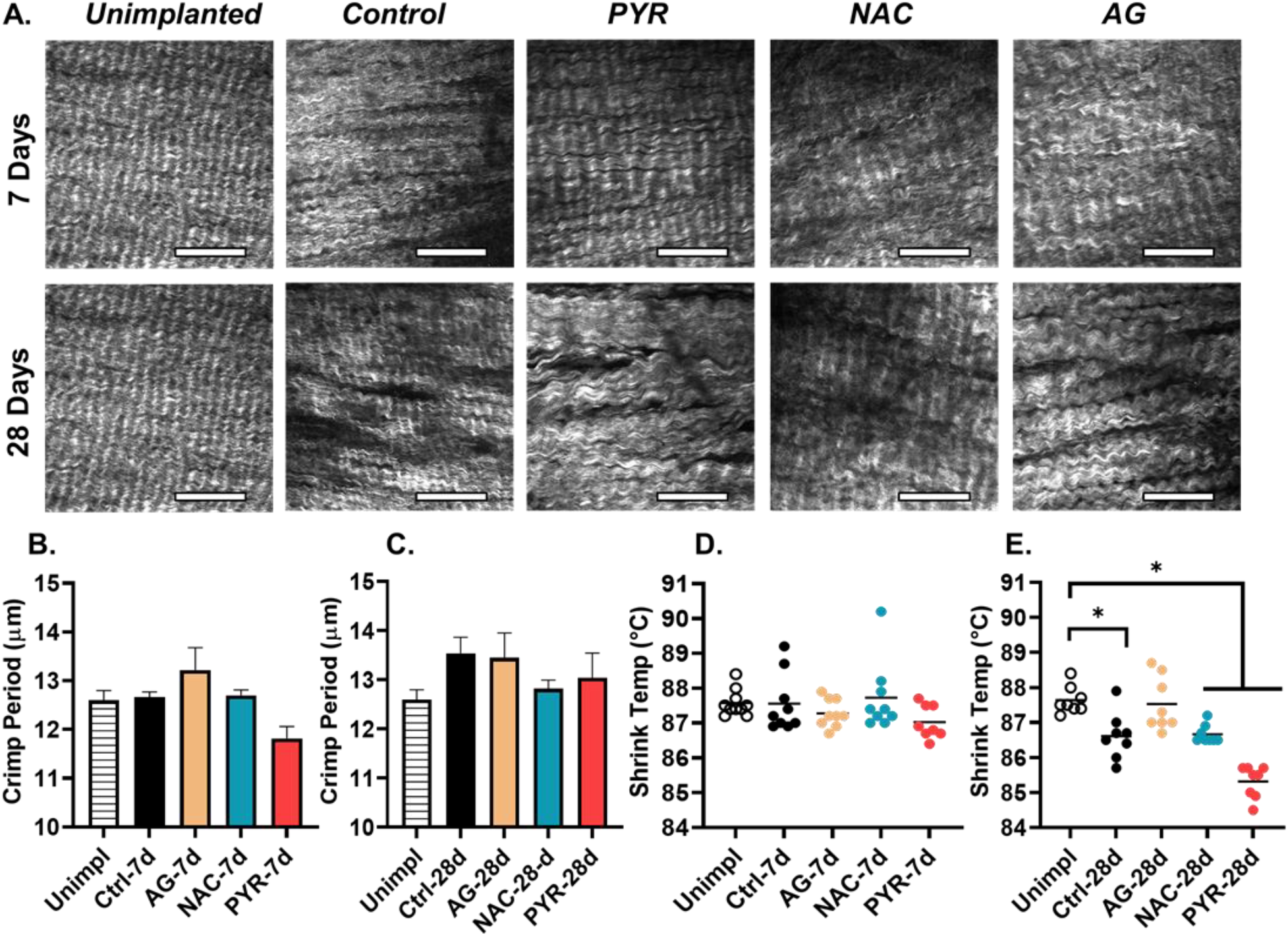
Second harmonic generation (SHG) imaging, crimp quantification, and shrink temperature values of glutaraldehyde-fixed bovine pericardium samples after 7- and 28-day implantation in juvenile rats. **A**. Representative SHG images of unimplanted control BP (*unimplanted*), implanted BP without drug administration (*control*) or after treatment with: aminoguanidine (*AG*), n-acetylcysteine (*NAC*), and pyridoxamine (*PYR*). Scale bars are 100 μm. **B**. Quantitative analysis of 7-day collagen crimp period based on microscopy images. **C**. Quantitative analysis of 28-day collagen crimp period based on microscopy images. **D**. Shrink temperature of 7-day explants. **E.** Shrink temperature of 28-day explants. Error bars indicate standard error. Significance was determined using Dunnett’s Method comparing to control (*p < 0.05).

Quantification of ST demonstrated no changes in control or drug groups after 7 days of implantation (Fig. 4D), while there was a significant decrease after 28 days in control (86.6±0.7°C), NAC (86.6±0.2°C), and PYR (85.3±0.4°C) (Fig. 4E). Thus, AGE inhibitors had a minimal effect on the collagen structure of implanted BP in vivo, while material properties were impacted after 28 days.

To understand the effects of inhibitors on glycation, albumin uptake, and calcification, samples were stained with specific antibodies for AGE and serum Albumin. Von Kossa staining and a quantitative calcium assay were used to assess the degree of calcification. There was minimal, yet positive staining using an αAGE antibody in control 7-day BP explants with a significant decrease in staining intensity from NAC and no change in AG or PYR compared to the unimplanted control (Supplementary Figure 2). Implantation for 28 days resulted in strong positive staining, which was not significantly impacted by AG or NAC (108±6%, 119±8% respectively), compared to the control explant (set as 100%) (Fig. 5A). Remarkably, PYR treatment resulted in almost 50% inhibition of staining intensity with 57±6% (Fig. 5A, PYR). Staining of 7-day explants using αAlbumin antibody resulted in a significant reduction in all of the AGE inhibitor groups, compared to the unimplanted control (55±1% for AG, 67±2% for NAC, and 42±4% for PYR) (Supplementary figure 3). After 28 days, only PYR had a significant mitigating effect on staining intensity with 30±5%, compared to the control (Fig. 5B). In contrast, NAC treatment resulted in a significant increase of staining intensity (135±3%), compared to the control (100%). No significant effect was observed in the AG group (107±5%).

**Figure 5.**
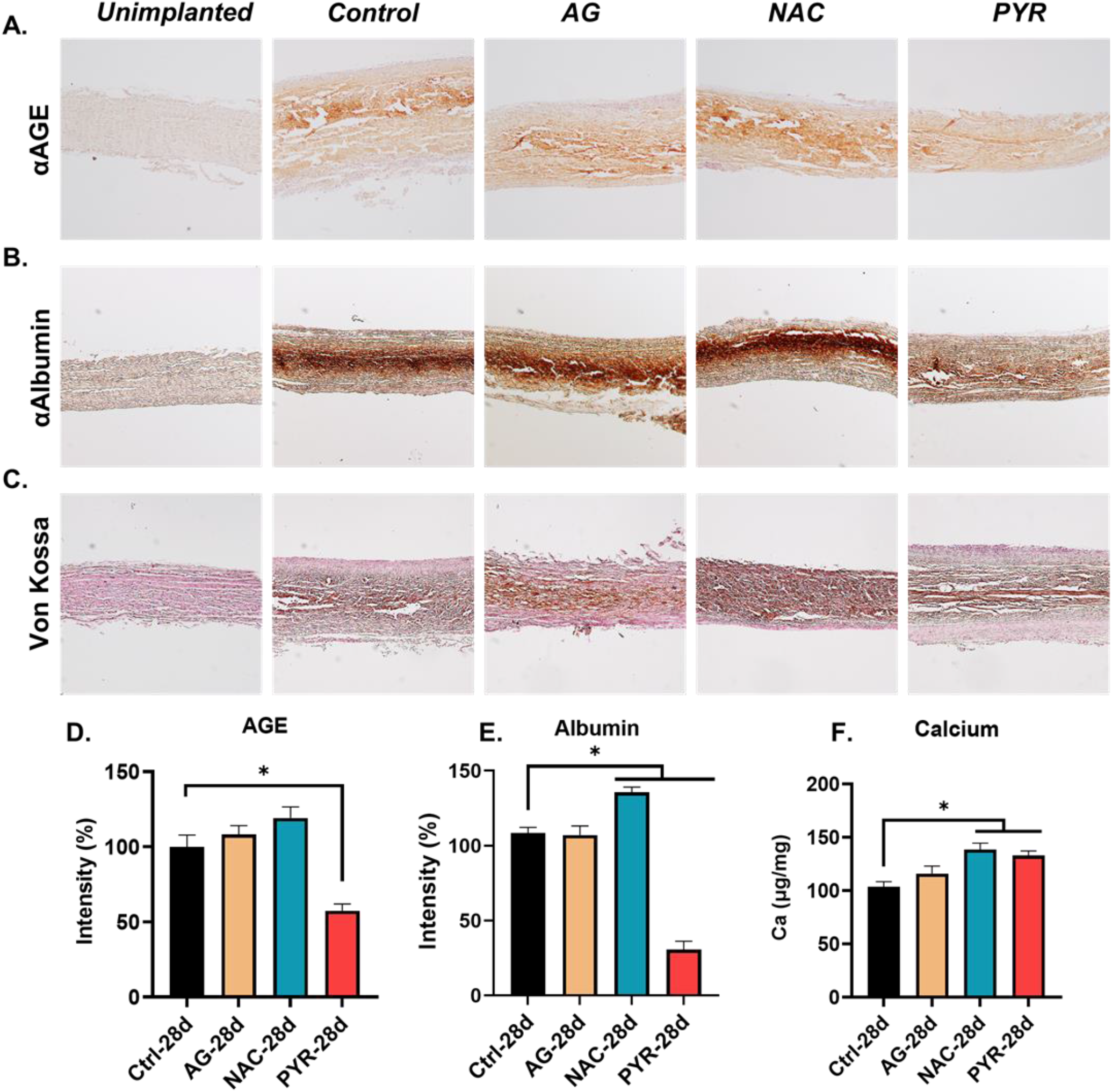
Analysis of rat subdermal explants after 28 days of implantation for AGE formation, Albumin uptake and calcification. **A**. Representative images of immunohistochemistry (IHC) staining for AGE of unimplanted BP (*unimplanted*) or after implantation without drugs (*control*), or after treatment with aminoguanidine (*AG*), n-acetylcysteine (*NAC*), and pyridoxamine (*PYR*) after 28 days of implantation. **B**. Representative images of IHC staining for Albumin after 28 days of implantation. **C.** Representative images of Von Kossa calcium specific staining after 28 days of implantation. **D.** Quantitative analysis of AGE IHC image staining intensity for 28-day samples. **E**. Quantitative analysis of Albumin image staining intensity for 28-day samples. **F**. Quantitative analysis of calcium content for 28-day samples. Significance was determined using Dunnett’s Method comparing to control (*p < 0.05).

Von Kossa calcium-specific staining was negative for all 7-day samples (Supplementary figure 4) and positive for all 28-day samples (Fig. 5C). The majority of calcium staining was concentrated in the middle region of the tissue, while NAC had a consistent intense staining across the tissue. Quantitative calcium analysis revealed no changes in calcification from AG while both NAC and PYR resulted in a significant increase in calcium staining. The observed changes pointed to the beneficial effect of PYR against pro-inflammatory AGE formation and albumin infiltration.

Serum levels of AGE metabolites were measured using enzyme-linked immunosorbent assay (ELISA) to study the impact of BP implantation and AGE inhibitors on the inflammatory processes in vivo. Rats treated with AGE inhibitors (AG, NAC, PYR) were compared with non-treated control rats. There was a significant increase in the levels of methylglyoxal measured 7 days after BP implantation from 6.2±1 μg/ml to 22±5 μg/ml (Fig. 6A, left), suggesting ongoing inflammation activity following the surgery. This increase was significantly mitigated by PYR resulting in 8.6±1 μg/ml, comparable to the levels measured in unoperated rats (Fig. 6A, left).

**Figure 6.**
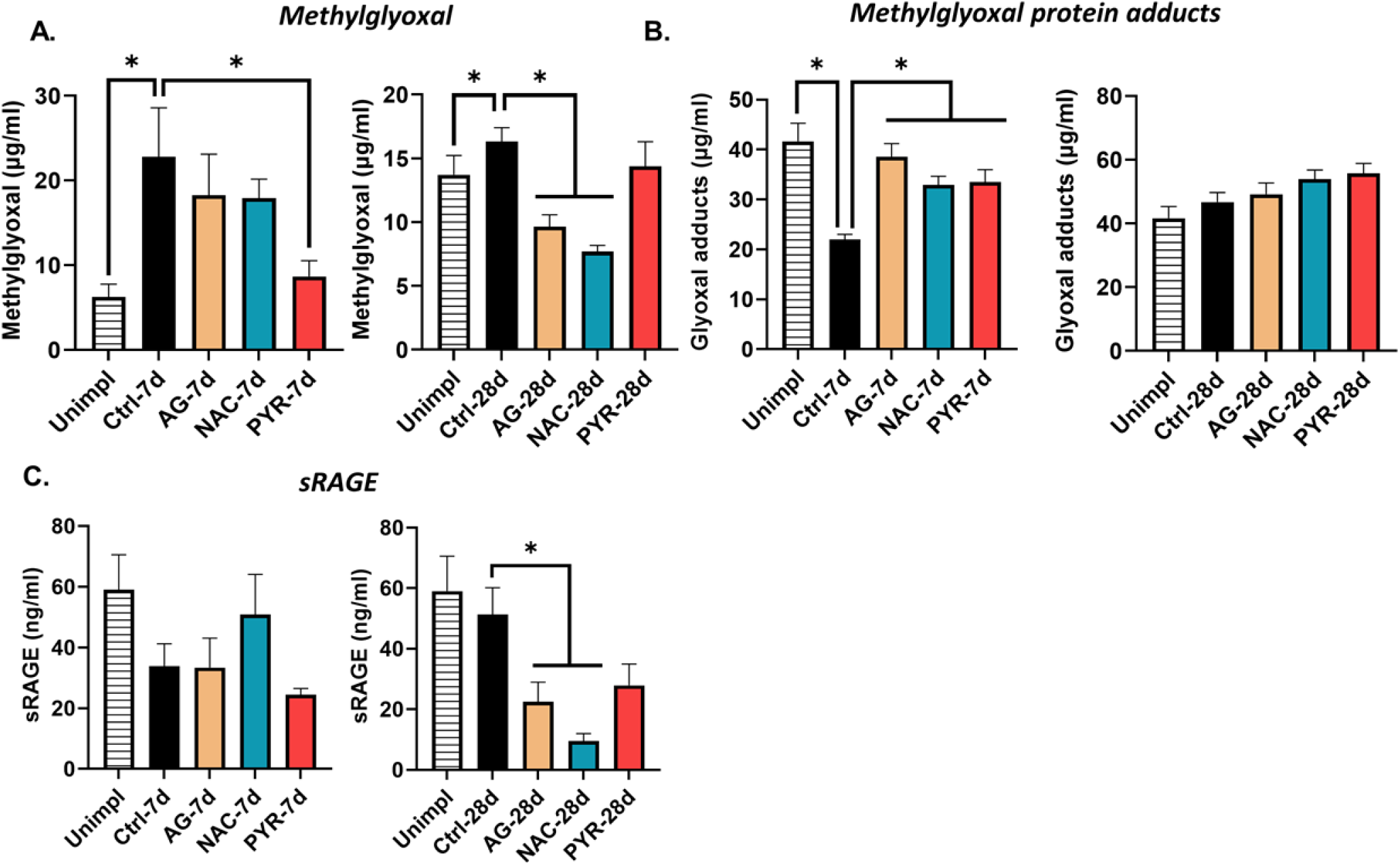
Serum levels of AGE metabolites in juvenile rats after 7 and 28 days of BP implantation with or without AGE inhibitors treatment measured by ELISA. **A**. Methylglyoxal serum levels unoperated control (*unimpl*), implanted BP without drug administration (*Control*) or after treatment with: aminoguanidine (*AG*), n-acetylcysteine (*NAC*), and pyridoxamine (*PYR*). **B**. Methylglyoxal protein adducts serum levels. **C**. Soluble receptor for AGE (sRAGE) levels. Significance was determined using Dunnett’s Method comparing to control (*p < 0.05).

No effect was measured in AG and NAC groups with 18.2±4 μg/ml and 17.9±2 μg/ml, respectively (Fig. 6A, left). After 28 days, methylglyoxal levels decreased to 16.3±1 μg/ml in control implanted animals and were similar in the unoperated control (13.7±1 μg/ml) (Fig. 6A, right). Similar values were measured in PYR treated animals (14.5±1 μg/ml), while there was a significant decrease in AG (10.1±0.8 μg/ml) and NAC (7.3±0.4 μg/ml) treated groups (Fig. 6A, right).

The levels of methylglyoxal protein adducts, a measure of AGE accumulation, were analyzed in the same samples (Fig. 6B). After 7 days of implantation, the levels showed a significant decrease from 40.3±4 μg/ml in the unoperated control to 21.1±1 μg/ml in the 7-day implanted control. Interestingly, all of the inhibitors showed significantly higher values than the 7-day implanted control (36.8±2 μg/ml for AG, 31.8±1 μg/ml for NAC, and 34.5±2 μg/ml for PYR). While there was an increase in all of the inhibitors after 7 days of implantation, compared to the untreated control, their levels were not different from ones in the unoperated control (Fig. 6B left).

Following 28 days of implantation, there was a two-fold increase in the levels of methylglyoxal protein adducts in the implanted control, compared to the 7-day unoperated control, resulting in 46±3 μg/ml, which was not significantly different from the 28-day unoperated control (41.4±4 μg/ml) (Fig. 6B right vs Fig. 6B left). No effect was measured from any of the inhibitors after 28 days (49.1±2 μg/ml for AG, 53.9±3 μg/ml for NAC, and 55.6±3 μg/ml for PYR).

The levels of sRAGE were measured to estimate the degree of inflammatory activity in serum. There was no significant effect on the levels of sRAGE in serum after 7 days of implantation (Fig. 6C left), while there was a significant decrease from AG (22.4±6 ng/ml) and NAC (9.5±2 ng/ml) after 28 days, compared to the implanted control (51.3±8 ng/ml) (Fig. 6C right).

All the inhibitors demonstrated glycation mitigation in vitro with mitigation of crimp collagen organization under accelerated glycation and reduction of radiolabeled glucose uptake by both BP and BSA. AGE inhibitors incubation resulted in a minimal yet significant change in BP shrink temperature. In vivo studies demonstrated that only PYR maintained significant anti-glycation effects towards BP implants, significantly reducing AGE formation and albumin infiltration. Calcification was not affected by AG or PYR, however incubation with NAC resulted in elevated calcification across the tissue. PYR treatment reduced the levels of circulating glyoxal in 7 days and maintained baseline levels of methylglyoxal.

## Discussion

The current study investigated, for the first time, the potential efficacy of three well established AGE inhibitors, AG, PYR, and NAC, for mitigating AGE-serum protein pathophysiology in BHV leaflets. It was recently demonstrated that AGE, serum protein deposition, and reactive oxygen species mediated processes are important contributors to SVD [3, 8, 11, 12]. Glycation occurs in all biological tissues during normal aging, resulting in pro-inflammatory epitope formation and tissue stiffening. Glycation is particularly accelerated in diabetes, resulting in earlier failure due to SVD of BHV in this patient population group [13]. Studies have demonstrated that BHV do not have a viable endothelial cell layer and therefore are particularly susceptible to glycation and serum protein infiltration [3].

AGE inhibitors were extensively studied to manage diabetic-related complications. It was shown that AG mitigated AGE formation in a diabetic rat model, as well as renal, retinal, neural, and vascular complications [14–17]. AG acted as a α, β-dicarbonyl scavenger and nitric oxide inhibitor, blocking formation of biologically active AGE. A vitamin B6 analogue known as PYR is another widely studied agent for mitigating AGE formation in vitro and in vivo. PYR has multiple mechanisms of mitigating AGE such as scavenging carbonyl species formed during Amadori product formation, binding metal ions that catalyze the reaction, and scavenging radicals [18–21]. Importantly, it was shown that PYR exhibits competitive inhibition of protein amino groups participating in reactions. Administration of PYR to diabetic rats reduced oxidative stress, decreased levels of circulating glyoxal, and mitigated AGE formation [5]. On the other hand, NAC was shown not only to scavenge reactive oxygen species and serve as a reducing agent, but also as an active cell metabolite inhibiting the NF-kB pathway [22].

In our study we explored the use of AG, NAC, and PYR against AGE formation in BHV using an established glycation model with in vitro endpoints of collagen structural integrity through SHG, shrink temperature and radiolabeled glucose uptake [3, 8]. All three inhibitors had a significant mitigating effect on radioactive glucose uptake in vitro by BP and BSA, while no significant effect was observed on the GLU-BSA co-incubation with BP, with the exception of NAC (Fig. 3). Furthermore, collagen image analysis of BP following incubation with BSA and NAC revealed a significant increase in BP crimp period (Fig. 1C). This data suggests the possible interaction between NAC and BSA in solution, possibly resulting in aggregate formation related to NAC sulfhydryl reactions, that can further infiltrate BP and disrupt collagen. Analysis of BP shrink temperature revealed a significant decrease in this parameter after AGE inhibitor co-incubations with glycation conditions, especially glucose. All inhibitors interact with BP to some extent, which would require further investigation (Fig. 2). The effect was less pronounced after 28-day subdermal implantation in rats, where only PYR and NAC had a significant effect on ST. There was a minimal effect of all drugs on the levels of sRAGE receptor with exception of AG and NAC. It was demonstrated that NAC has a mitigating effect on sRAGE formation [6]. Despite the beneficial effect on this metabolic marker, NAC showed unfavorable effects on albumin accumulation and calcification, resulting in acceleration of these processes.

The results of in vitro and in vivo experiments demonstrated consistent anti-glycation effects of PYR while maintaining the integrity of the collagen microstructure (Fig. 1, Fig. 2, Fig. 5). Surprisingly, there was superior mitigation of uptake of a major protein susceptible to glycation, serum albumin, by PYR in rat subdermal implants (Fig. 5). This could be due to a possible interaction between PYR and BP aldehyde groups, resulting in tissue modification with PYR. The results of shrink temperature analyses, a measure of collagen crosslinking, also show a significant decrease after PYR-BP co-incubation, pointing to the possible modification.

The in vivo studies, analyzing serum biomarkers, demonstrated a significant reduction of circulating methylglyoxal in rats after 7 days of BP implantation and PYR administration. While there was no significant decrease of methylglyoxal after 28 days, the levels were similar to those seen in control unoperated animals, and were consistent with previously published data reporting no long-term PYR effect on methylglyoxal levels in non-diabetic rats.

We can conclude that PYR is a promising anti-AGE agent with a potential application for patients with BHV, offsetting SVD through the mitigation of AGE-related complications and the preservation of valve collagen microstructure. This work offers novel insights for SVD preventative treatment, providing a possible solution for an unmet clinical need.

## Acknowledgements

This work was supported by the following sources: NIH grants HL143008 (RJL and GF) and HL007915 (RJL, AZ and CR), Congenital Heart Defect Coalition (AZ), The Pediatric Heart Valve Center of the Children’s Hospital of Philadelphia (RJL), The Kibel Fund for Aortic Valve Research (to GF and RJL), The Valley Hospital Foundation ‘Marjorie C Bunnel’ charitable fund (to GF), the Andrew Sabin Family Foundation Research Laboratory (GF) and both Erin’s Fund and the William J Rashkind Endowment of the Children’s Hospital of Philadelphia (to RJL).The authors thank Susan Kerns for manuscript preparation and submission.

