## Supplemental Figures for "Inhibition of advanced glycation end product formation and serum protein infiltration in bioprosthetic heart valve leaflets: Investigations of anti-glycation agents"

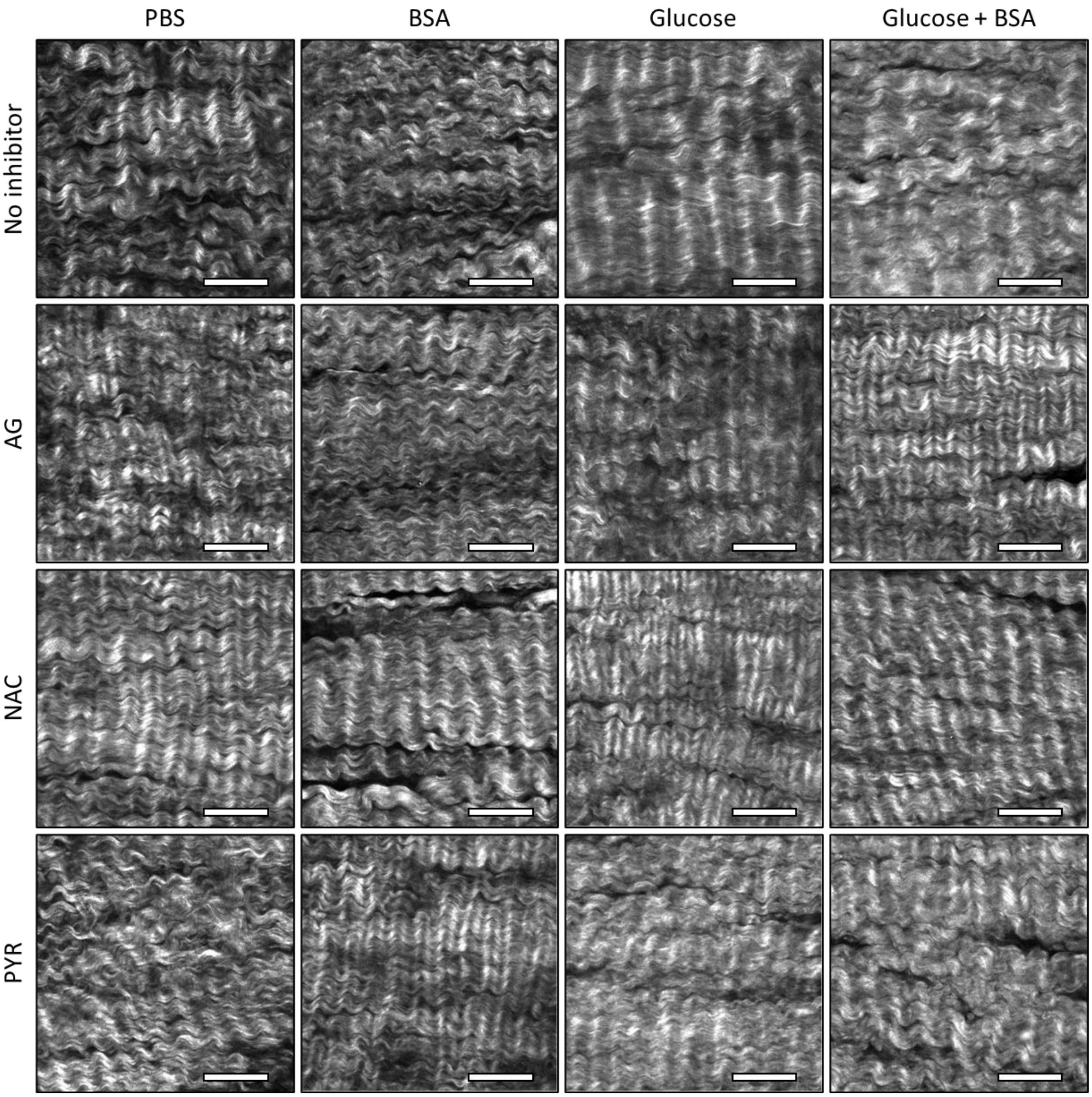


**Supplementary Figure 1. Representative two-photon microscopy images of glutaraldehyde-fixed bovine pericardium following 28 days of exposure to the following glycation inhibitors: none (control), aminoguanidine (AG 10mM), n-acetylcysteine (NAC, 5mM), and pyridoxamine (PYR, 50mM).** The inhibitors were dissolved in: PBS, PBS with bovine serum albumin (BSA, 5%), PBS with glucose (100mM), PBS with glucose (100mM) plus BSA (5%). Scale bar is 50µm.

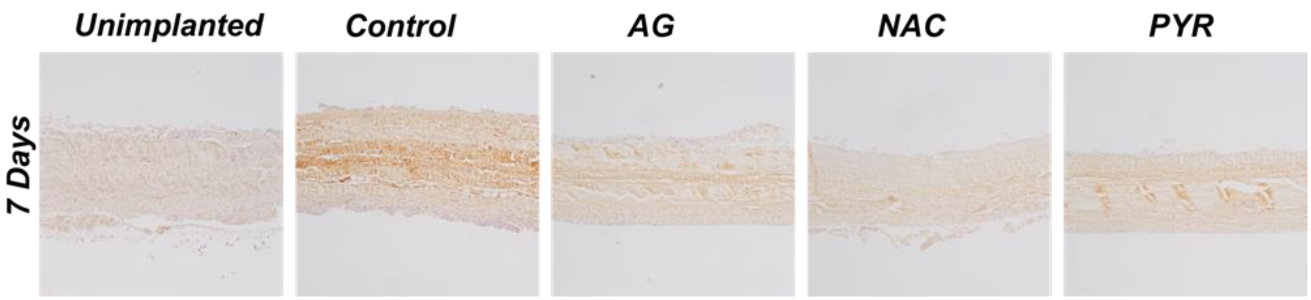


**Supplementary Figure 2. Immunohistochemical analysis of rat subdermal explants after 28 days of implantation for AGE formation.** Representative images of immunohistochemistry staining for AGE of unimplanted BP (unimplanted) or after implantation without drugs (control), or after treatment with aminoguanidine (AG), n-acetylcysteine (NAC), and pyridoxamine (PYR).

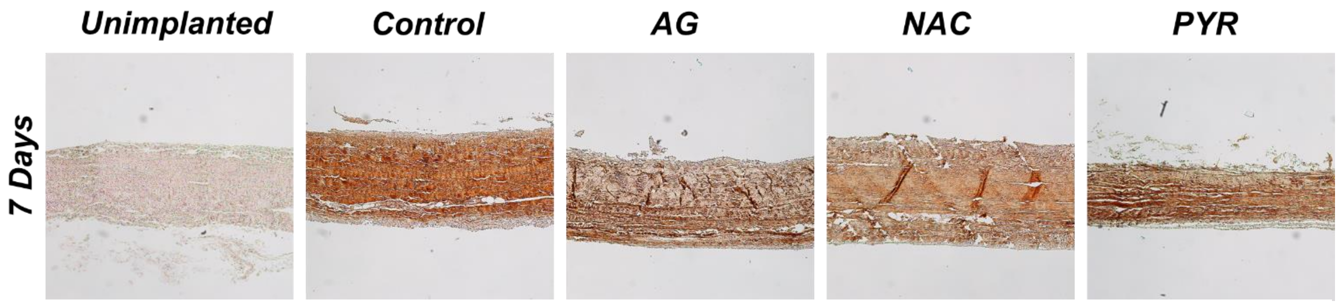


**Supplementary Figure 3. Immunohistochemical analysis of rat subdermal explants after 28 days of implantation for serum albumin albumin uptake.** Representative images of immunohistochemistry staining for serum albumin of unimplanted BP (unimplanted) or after implantation without drugs (control), or after treatment with aminoguanidine (AG), n-acetylcysteine (NAC), and pyridoxamine (PYR).

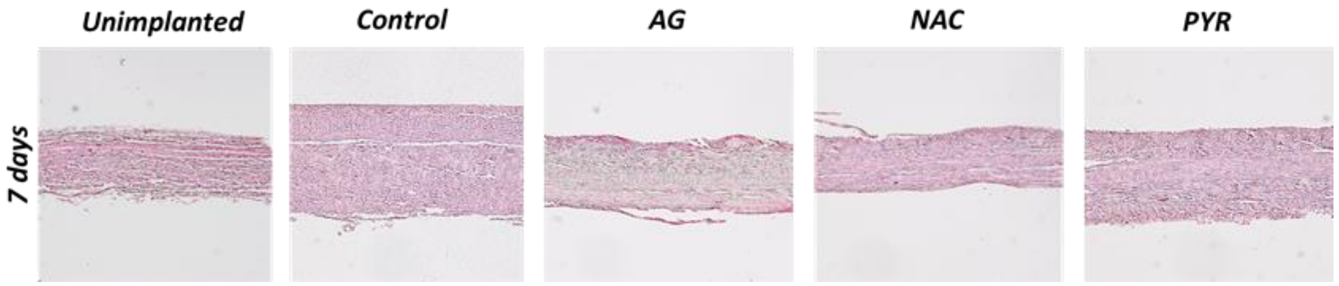


**Supplementary Figure 4.** Representative images of Von Kossa calcium specific staining after 7 days of implantation of unimplanted BP (unimplanted) or after implantation without drugs (control), or after treatment with aminoguanidine (AG), n-acetylcysteine (NAC), and pyridoxamine (PYR).
